# Heavily and Fully Modified RNAs Guide Efficient SpyCas9-Mediated Genome Editing

**DOI:** 10.1101/290999

**Authors:** Aamir Mir, Julia F. Alterman, Matthew R. Hassler, Alexandre J. Debacker, Edward Hudgens, Dimas Echeverria, Michael H. Brodsky, Anastasia Khvorova, Jonathan K. Watts, Erik J. Sontheimer

**Author notes:** Correspondence (A.K.); (J.K.W.), (E.J.S.).

## Abstract

RNA-based drugs depend on chemical modifications to increase potency and nuclease stability, and to decrease immunogenicity *in vivo*. Chemical modification will likely improve the guide RNAs involved in CRISPR-Cas9-based therapeutics as well. Cas9 orthologs are RNA-guided microbial effectors that cleave DNA. No studies have yet explored chemical modification at all positions of the crRNA guide and tracrRNA cofactor. Here, we have identified several heavily-modified versions of crRNA and tracrRNA that are more potent than their unmodified counterparts. In addition, we describe fully chemically modified crRNAs and tracrRNAs (containing no 2’-OH groups) that are functional in human cells. These designs demonstrate a significant breakthrough for Cas9-based therapeutics since heavily modified RNAs tend to be more stable *in vivo* (thus increasing potency). We anticipate that our designs will improve the use of Cas9 via RNP and mRNA delivery for *in vivo* and *ex vivo* purposes.

---

CRISPR RNA-guided genome engineering has revolutionized research into human genetic disease and many other aspects of biology. Numerous CRISPR-based *in vivo* or *ex vivo* genome editing therapies are nearing clinical trials. At the heart of this revolution are the microbial effector proteins found in class II CRISPR-Cas systems^1^ such as Cas9 (type II) and Cas12a/Cpf1 (type V).^2-4^

The most widely used genome editing tool is the type II-A Cas9 from *Streptococcus pyogenes* strain SF370 (SpyCas9)^2^. Cas9 forms a ribonucleoprotein (RNP) complex with a CRISPR RNA (crRNA) and a trans-activating crRNA (tracrRNA) for efficient DNA cleavage both in bacteria and eukaryotes (Figure 1A). The crRNA contains a guide sequence that directs the Cas9 RNP to a specific locus via base-pairing with the target DNA to form an R-loop. This process requires the prior recognition of a protospacer adjacent motif (PAM), which for SpyCas9 is NGG. R-loop formation activates the His-Asn-His (HNH) and RuvC-like endonuclease domains that cleave the target strand and the non-target strand of the DNA, respectively, resulting in a double-strand break (DSB).

**Figure 1:**
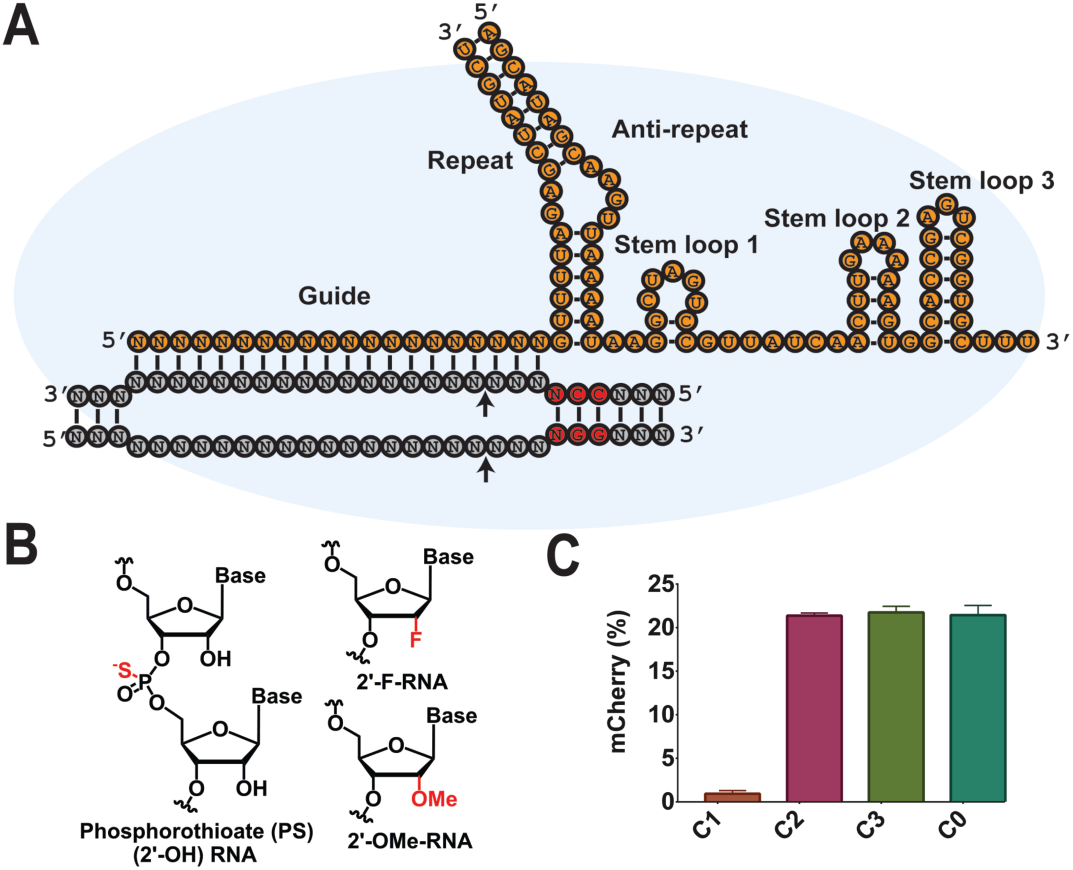
Initial screening of chemical modifications in the crRNA. **A**. Schematic of Cas9 RNP paired with target DNA. The secondary structure elements of crRNA and tracrRNA are labeled. RNA is shown in orange, whereas DNA is in grey. The PAM sequence is highlighted red and cleavage sites are marked with arrows. **B**. Chemical modifications used in this study. **C**. Bar graph showing mCherry-positive cells after nucleofection of HEK293T-TLR cells with RNPs that included the indicated crRNAs and an unmodified tracrRNA. Error bars represent standard deviation (SD) resulting from at least three biological replicates.

For mammalian applications, Cas9 and its guide RNAs can be expressed from DNA (e.g. a viral vector), RNA (e.g. Cas9 mRNA plus guide RNAs in a lipid nanoparticle), or introduced as an RNP. Viral delivery of Cas9 results in efficient editing, but can be problematic because long-term expression of Cas9 and its guides can result in off-target editing, and viral vectors can elicit strong host immune responses.^5^ RNA and RNP delivery platforms of Cas9 are suitable alternatives to viral vectors for many applications and have recently been shown to be effective genome editing tools *in vivo*.^6,7^ RNP delivery of Cas9 also bypasses the requirement for Cas9 expression, leading to faster editing. Furthermore, Cas9 delivered as mRNA or RNP exists only transiently in cells and therefore exhibits reduced off-target editing. For instance, Cas9 RNPs were successfully used to correct hypertrophic cardiomyopathy (HCM) in human embryos without measurable off-target effects.^8^

The versatility of Cas9 for genome editing derives from its RNA-guided nature. The crRNA of SpyCas9 used in this study consists of a 20-nt guide region followed by a 16-nt repeat region (Figure 1A). The tracrRNA consists of an anti-repeat region that pairs with the crRNA, and also includes three stem-loops. All of these secondary structure elements are required for efficient editing in mammalian systems.^9^ However, unmodified RNAs are subject to rapid degradation in circulation and within cells.^10,11^ Therefore, it is highly desirable to chemically protect RNAs for efficient genomic editing in hard-to-transfect cells and *in vivo*. Thus, it has been previously reported that chemical modifications in the crRNA and tracrRNA enhance stability and editing efficiency *in vivo* and *ex vivo*.^6,7,11-13^ Chemical modifications including 2’-*O*-methyl (2’-OMe), phosphorothioate (PS), 2’-*O*-methyl thioPACE (MSP), 2’-fluoro RNA (2’-F-RNA) and constrained ethyl (S-cEt) have previously been employed to synthesize crRNA and tracrRNA.^6,11,12^ The modified RNAs not only improved Cas9 efficacy, but in some instances also improved specificity.^11,14^ Modifications were either based on the crystal structures of Cas9 or limited to the ends of RNAs, and the guides were not modified extensively. Nonetheless, heavily or fully modified RNAs may have advantages *in vivo*.^10^ Modified siRNAs and ASOs substantially improve stability and potency, and can also reduce off-target effects. Furthermore, extensively modified RNAs can prevent innate immune responses.^15^

In the present study, we sought to extensively modify the crRNA and tracrRNA while retaining the efficacy of SpyCas9-based genome editing in cultured human cells. We used structure-guided and systematic approaches to introduce 2’-OMe-RNA, 2’-F-RNA and PS modifications (Figure 1B) throughout guide RNAs (Table S1). Our strategy yielded active RNP complexes with both extensively and fully modified versions of crRNAs and tracrRNAs.

Crystal structures of SpyCas9 have been solved as the RNP alone or bound to one or both strands of target DNA.^16-19^ These structures provide detailed information regarding the interactions between the Cas9 protein and crRNA:tracrRNA complex. We used these structures to identify sites where Cas9 protein makes no contacts with the crRNA or tracrRNA. Thus, in our initial screen, 2’-OMe modifications were introduced at guide positions 7-10 and 20 (**C2**, Figure 1C). Similarly, positions 21 and 27-36 in the crRNA repeat region were also modified using 2’-OMe. To improve nuclease stability, PS modifications were also introduced at the 5’ end of the crRNA, yielding the **C3** design (Figure 1C and Table 1). In parallel, we tested a crRNA that was more aggressively modified to leave only nine nucleotides (nt) unprotected (**C1**). Similarly, 2’-OMe modifications were also introduced into the tracrRNA at all positions where no protein contact with the RNA is observed. This gave rise to **T1** that is 50% chemically modified (Figure 2 and Table 2).

**Table 1:**
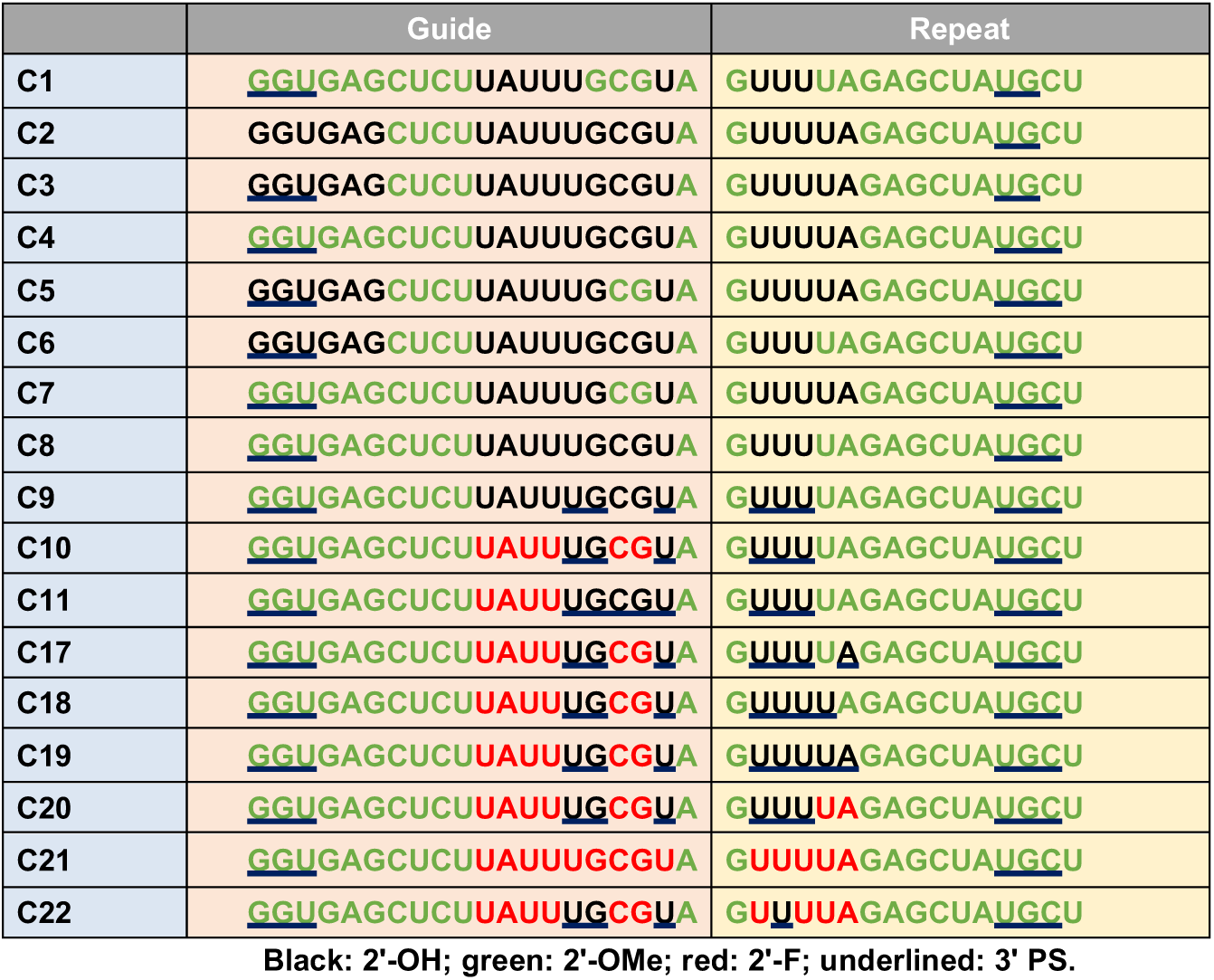
Chemically modified crRNAs used in this study

**Table 2:**
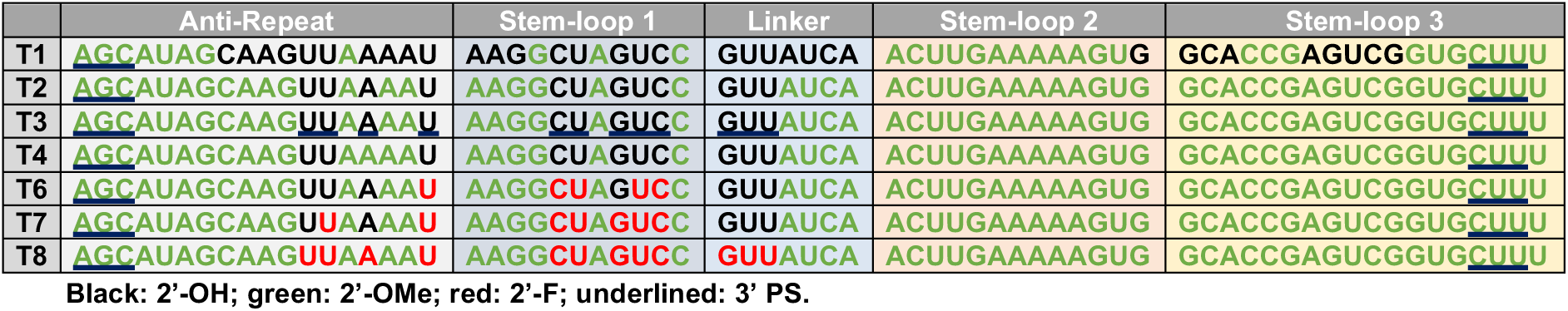
The tracrRNA sequences used in this study

**Figure 2:**
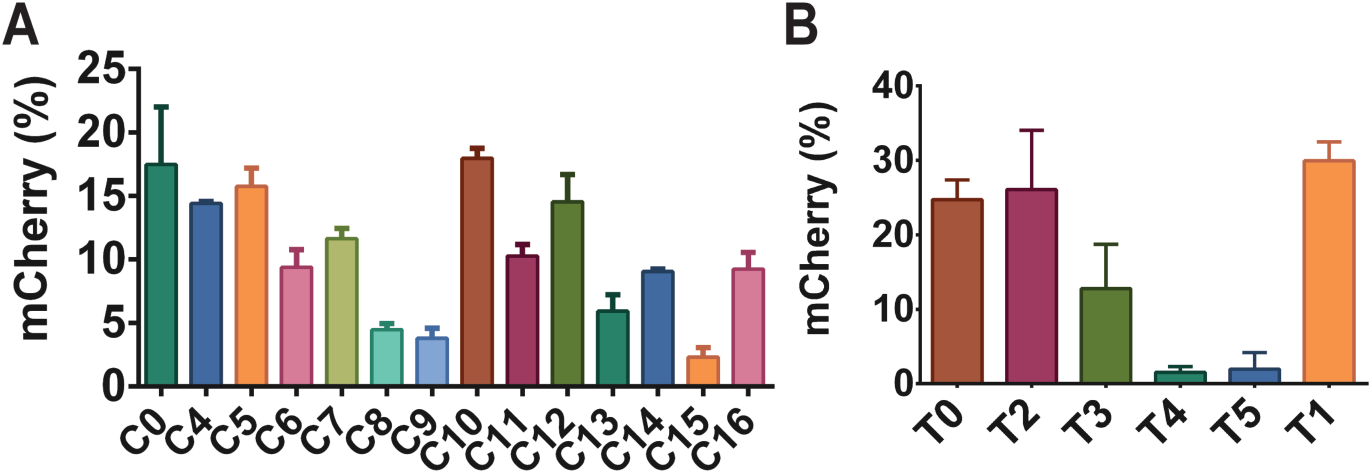
Second round of chemical optimization of crRNA (**A**) and tracrRNA (**B**). Each crRNA was tested with the unmodified tracrRNA **T0**, whereas each tracrRNA was tested with the unmodified crRNA **C0**. The bar graphs show the percent of cells expressing mCherry, ± SD. Each RNA was tested in triplicate.

The crRNAs and tracrRNAs were tested in a HEK293T cell line stably expressing the traffic light reporter (TLR) system.^20^ The HEK293T-TLR cells were nucleofected (Neon Transfection System) with an *in vitro*-reconstituted RNP complex of recombinant 3xNLS-SpyCas9, crRNA and tracrRNA. The nucleofected cells were analyzed by flow cytometry for mCherry-positive cells, which reports on a subset of non-homologous end-joining DSB repair events.^20^ As shown in Figure 1C, modified crRNAs 2 and 3 retain complete activity relative to the unmodified crRNA **C0**, suggesting that the modifications introduced in crRNAs 2 and 3 are well tolerated by Cas9. Lipid-based delivery of Cas9 RNP complex showed that **C3** was even more efficacious than **C0** and **C2** (Figure S1), which demonstrates the importance of PS linkages at the 5’ terminus of the crRNA. Similarly, **T1** did not hinder Cas9 activity. On the other hand, the extra modifications introduced in **C1** almost completely abolished Cas9 activity in cells. We reasoned that the 2’-OMe modifications (especially at positions 16-18 in the crRNA) are most likely to compromise Cas9 RNP activity since nt at position 16 and 18 were shown to make base-specific contacts with Arg447 and Arg71.^16^ The 2’-OH of G16 in the TLR crRNA is also predicted to make a hydrogen bond with Arg447. We chose **C3** and **T1** as a basis for further optimization.

In the second round of crRNA modification, we introduced additional 2’-OMe modifications into the first 6 nt of **C3** to yield **C4** (Figure 2). In another design, 2’-OMe modifications were incorporated at positions 17 and 18 (**C5**). G16 was left unmodified because it makes base- and backbone-specific contacts with Cas9 and likely contributed to the low efficacy of **C1**. Recently, others have also observed similar constraints at position 16.^6^ In **C6**, the importance of 2’-OH groups at positions 25 and 26 was tested. The 2’-OH of these nts contacts the protein in the crystal structure; however, they do not pair with the target DNA, and 2’-OMe substitution at these positions may therefore be more tolerable. **C7** and **C8** were identical to **C5** and **C6**, respectively, except that they also contained 2’-OMe modifications in the first six positions. All of these crRNAs (**C4**-**C8**) were designed to identify modifications responsible for the lower activity of **C1** relative to **C3**.

As shown in Figures 2 and S2, **C4-C7** retain almost the same efficacy as **C0**, but **C8** activity was strongly reduced. These results indicated that nts at positions 1-6 and 17-18 tolerate 2’-OH substitutions. 2’-OMe modifications at positions 25 and 26 were tolerated in **C6** but not in **C8**. We had also synthesized a version of **C8** that contained PS linkages at several unprotected positions including 15-16, 19 and 21-23 (**C9**). This design also exhibited reduced editing efficiency by Cas9. When tested for DNA cleavage activity *in vitro*, **C8** and **C9** were fully active even at low RNP concentrations (Figure S3). These results suggest structural perturbations in **C8** and **C9** that are particularly acute under intracellular conditions.

We also incorporated 2’-F-RNAs in this round of optimization since they can increase thermal and nuclease stability of RNA:RNA or RNA:DNA duplexes, and they also interfere minimally with C3’-endo sugar puckering.^21,22^ 2’-F may be better tolerated than 2’-OMe at positions where the 2’-OH is important for RNA:DNA duplex stability. For these reasons, we synthesized two crRNAs based on **C9** but with 2’-F modifications at positions 11-14 and/or 17-18 (**C10-C11**). These modifications rescued some of **C9**’s diminished activity. In fact, **C10 (**which contained 2’-F substitutions at positions 11-14 and 17-18) performed better than **C11**, in which positions 17-18 were unmodified. Our results suggest that 2’-F substitutions can compensate for lost efficacy resulting from high 2’-OMe content. It is especially noteworthy that **C10** retains the same activity as the unmodified **C0** but contains at least one backbone modification at every single phosphodiester linkage. This represents a significant breakthrough for Cas9-based therapeutics because **C10** has great potential to provide increased stability, and therefore more efficient editing, *in vivo*.

We also carried out a second round of tracrRNA optimization. **T1** was further modified by introducing 2’-OMe substitutions at most positions where the 2’-OH groups do not make crystal contacts with the protein. In addition, some nts that interact with Cas9 were also modified, given that the crRNA tolerated substitutions at many such positions. This approach produced tracrRNAs **T2**-**T5**, which contain modifications in at least 55 out of 67 nt. A15 is the only position that differs between **T2** and **T4** whereas **T3** contains additional stabilizing PS linkages at unprotected positions relative to **T2**. These tracrRNAs were tested in HEK293T-TLR cells, and the majority of 2’-OMe chemical modifications were tolerated by the tracrRNA except at position A15 (Figure 2). In the crystal structure, the 2’-OH of A15 interacts with Ser104. The best-performing tracrRNA from this round was **T2**, which contains 12 unmodified positions. Furthermore, the inclusion of PS linkages at these 12 positions reduced but did not abolish activity. This design (**T3**) contains at least one chemical modification at every position (either a PS or ribose modification). This also represents an important advance for therapeutic applications of Cas9.

The mCherry signal only results from indels producing a +1 frameshift, and therefore underestimates true editing efficiencies. To ensure that crRNA:tracrRNA combinations do not yield false negatives by favoring TLR indels that are out of the mCherry reading frame, we also carried out Tracking of Indels by Decomposition (TIDE) analysis to analyze overall editing efficiencies. As shown in Figure S2, editing efficiencies measured using TIDE correlate well with the mCherry signal.

We also explored whether addition of terminal modifications such as fluorophores, *N*-Acetylgalactosamine (GalNAc), or Cholesterol-Triethylene glycol (TEGChol) are tolerated by the crRNA and the tracrRNA. Such modifications can be useful for microscopy, and for monitoring cellular or tissue-specific RNA uptake. We introduced 5’-Cy3 modifications on crRNAs **C10** and **C11** to yield **C12** and **C13**, respectively (Table S1). We also covalently attached TegChol or GalNAc to the 3’ end of **C12** or **C13** to obtain **C14** and **C15**, respectively. Most crRNA modifications were tolerated on both ends, though some loss of function was observed with **C13, C14** and **C16** (Figure S2). In contrast, **C15** was essentially inactive. **T5** containing a 3’-TegChol was also nonfunctional, not surprisingly given the 2’-OMe substitution at A15.

We built upon the best-performing individual chemical configurations (**C10** and **T2**) to attempt to define combined crRNA:tracrRNA modification patterns that are compatible with SpyCas9 RNP function. Because crRNA 2’-F substitutions were largely tolerated (Figure 2), and in some cases even compensated for the loss of efficacy caused by 2’-OMe substitutions, we added several 2’-F modifications to **C10** and **T2**. In addition, because we had observed that simultaneous 2’-OMe modification at positions 25 and 26 negatively affected efficacy in some cases (e.g. **C8**), we tested these two positions for their sensitivities to 2’-F or individual 2’-OMe substitutions. We also incorporated additional 2’-F modifications in the tracrRNAs. In positions where the nucleobases interact with Cas9, we took two approaches to modification. While we suspected that protein-interacting sites would be less tolerant of modification, it was difficult to predict whether steric constraints or charge interactions were more important. To address this issue, we synthesized three different tracrRNAs: one where all protein interacting sites were left as 2’-OH (**T6**), another where all were converted to 2’-F (**T8**), and another where only the nucleobases that interact with nonpolar amino acids were converted to 2’-F (**T7**). Using this systematic approach, crRNAs **C17-C22** and tracrRNAs **T6-T8** were synthesized and tested (Figure 3A).

**Figure 3:**
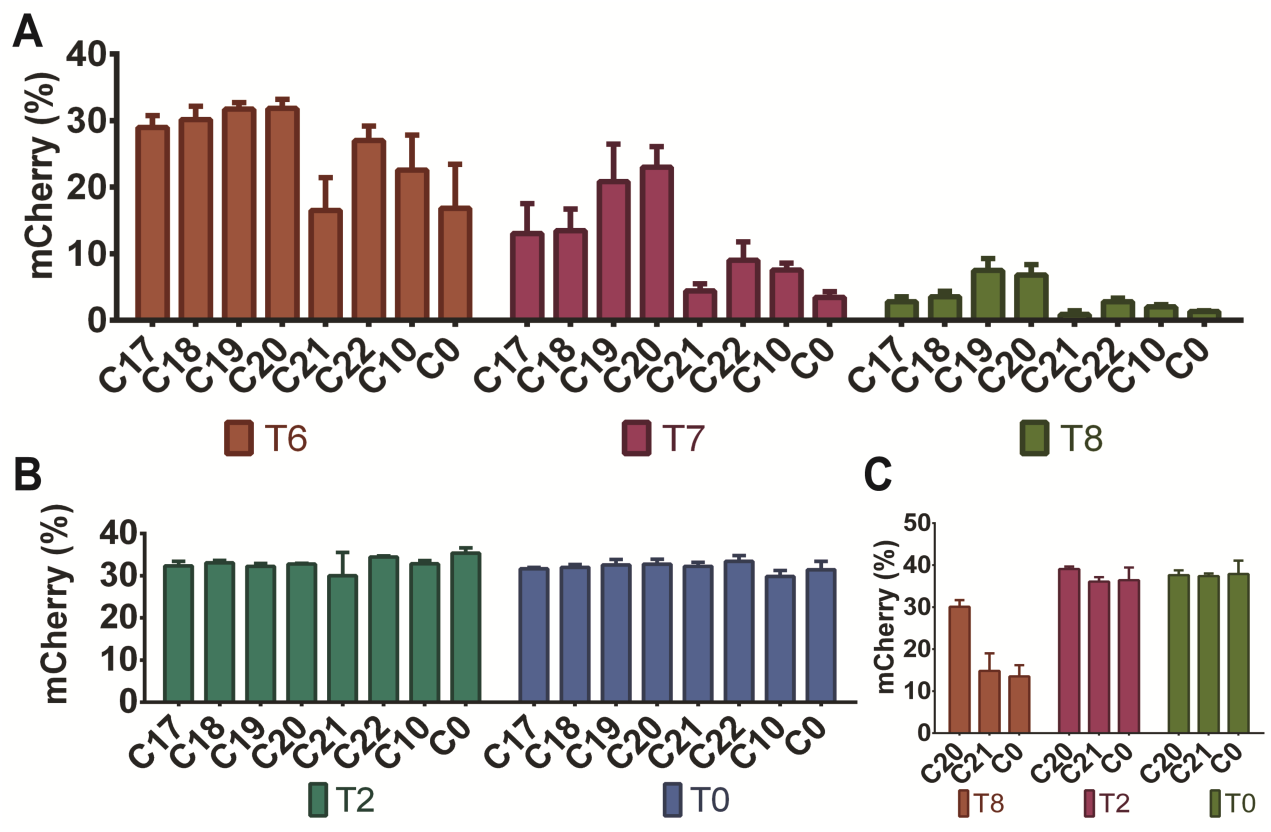
Cas9 tolerates heavily- and fully-modified crRNA:tracrRNA. **A** & **B**. Each crRNA was tested with tracrRNAs **T0, T2**, and **T6-T8** using 20 pmol of Cas9 RNP. **C**. HEK293T-TLR cells were also nucleofected with 100 pmol of the indicated RNPs to test whether heavily modified RNAs regain functionality at higher doses. Error bars show ± SD of three biological replicates.

When **C17-C22** were used with either **T2** or the **T0** control (20 pmol RNP), all showed comparable efficacy as the **C0** and **C10** crRNAs (Figure 3B). This includes the fully modified **C21** that is either 2’-F- or 2’-OMe-substituted at every position. To our knowledge, this is the first time a completely modified and fully functional crRNA has been reported. **C21** loses some efficacy when combined with **T6-T8**, and is also less potent than **C0** when lower (3 pmole) doses of RNP are delivered (Figure S4). These losses may be due to compromised base pairing between the heavily modified repeat:anti-repeat duplexes. Across all tracrRNAs tested, **C20** exhibits the highest editing efficiency. In addition, at 3 pmol RNP, **C20** is more potent than unmodified **C0**, suggesting enhanced stability in cells (Figure S4). Although **C20** includes six ribose sugars, each is adjacent to a PS modification, leaving no unmodified linkages in the crRNA.

Among **T6**-**T8**, the best-performing tracrRNA was **T6**, especially with modified crRNAs including **C20**. The fully-modified tracrRNA (**T8**) compromised the potency of all crRNAs tested, but retains some function (∼5% editing with 20 pmol RNP) with **C19** and **C20** (Figure 3B). To test whether the **T8** activity improves at higher doses, we nucleofected cells with 100 pmol Cas9 RNP. We found that by using a higher amount of Cas9 RNP, the editing efficiency of **T8** in combination with **C0** or **C20** is rescued to the same level as observed using 20 pmol of Cas9 RNP with **C0:T0** (Figure 3). Furthermore, at higher doses, the efficacy of **C20**:**T8** is almost as high as that of **C20**:**T0**. Lastly, the editing efficiency of the fully-modified pair (**C21**:**T8**) is within ∼2-fold of the unmodified (**C0**:**T0**) crRNA:tracrRNA pair. To our knowledge, this is the first demonstration of efficient editing activity with a fully-modified crRNA:tracrRNA combination. While the editing efficiency is not as high as that of the unmodified RNAs in cells, the increased serum stability afforded by the fully chemically optimized **C21**:**T8** combination (Figure S5) would likely provide significant benefits *in vivo*, as observed for fully modified siRNAs and ASOs.

To verify that our crRNA designs are compatible with different guide sequences, including those targeting endogenous human genes, we tested the **C10, C20** and **C21** designs targeting the huntingtin (*HTT*), human hemoglobin β (*HBB*), and Vascular Endothelial Growth Factor A (*VEGFA*) genes.^14,23^ *VEGFA* and *HBB* target sites were chosen for their therapeutic potential as well as the fact that they have been previously validated for genome editing. The *HTT* site, on the other hand, is a potential polymorphic target for Huntington’s disease treatment. As shown in Figure 4A & 4B, *HTT-***C10** and *HTT* **-C20** performed as well as the minimally modified *HTT*-**C0** when paired with **T2** and **T0. T6** and **T7** are more efficacious with the modified **C10** compared to minimally modified **C0**. The fully modified *HTT*-**C21** performed as well as the *HTT-* **C0** when tested with **T2**. However, similar to the TLR target site, some loss of potency is observed with the fully modified **T8**. However, **T8** did support editing with efficiencies comparable to **T0** when paired with **C20**. Similar results were obtained at the *HBB* and *VEGFA* target sites (Figure 4C & 4D): our potent RNA designs (**C20:T2**) performed as well as the minimally modified designs, and the fully modified dual guides exhibited some loss in potency. Furthermore, nucleofections performed using 3 pmol of RNP suggested that **C10** and **C20** may be more efficacious (but never less efficacious) than the unmodified crRNA, similar to what was observed in Figure S4, but this effect seemed to vary between target sites (Figure S8). **C20** also showed higher potency compared to **C0** when tested in human embryonic stem cells (hESC) (Figure 4E). In hESC the highest potency was achieved using the heavily modified combination **C20:T2**. Furthermore, the fully modified crRNA **C21** was just as efficacious as the minimally modified **C0**. We also examined editing in HEK293T cells at the top off-target site for both *HBB* and *VEGFA*, as validated previously.^14,23^ The modified crRNAs do not significantly affect off-target editing, though the fully modified **C21:T8** may provide slight specificity improvements compared to the less heavily modified designs (Figure S9). Collectively these results demonstrate that our modified crRNA designs can be applied to endogenous target sites.

**Figure 4:**
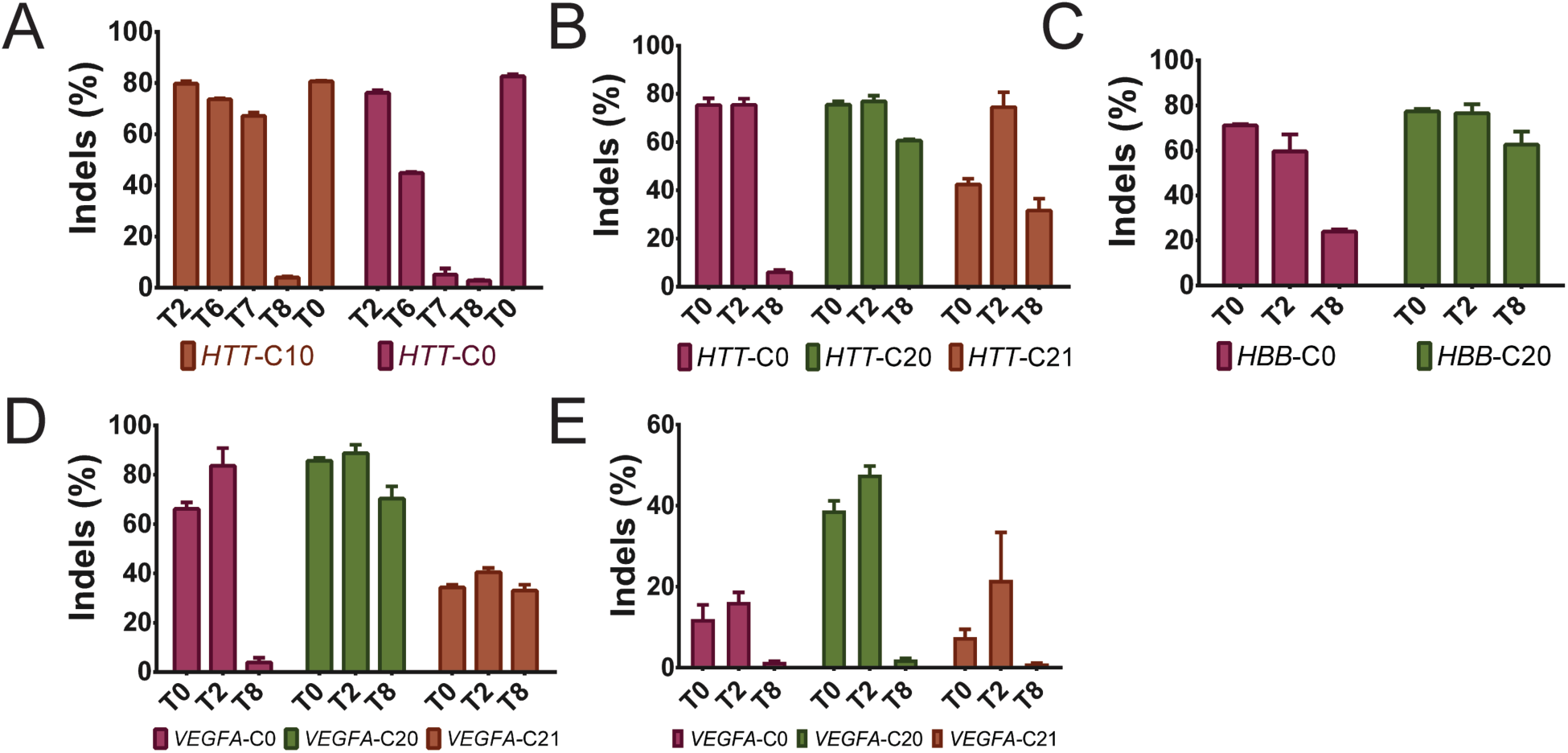
Targeting endogenous genes with modified RNAs. **A**. The **C10** guide design targeting *HTT* exon 50 was tested using 20 pmol of RNP along with **T2** and **T6-T8** in HEK293T cells. **B**. *HTT*-**C20** and *HTT*-**C21** designs were tested using 80 pmol of Cas9 RNP with the indicated tracrRNAs. **C**. The therapeutically relevant *HBB* locus was targeted using a previously validated guide sequence incorporated into the **C20** design. **D-E**. *VEGFA*-targeting crRNAs **C20** and **C21** were tested using the indicated tracrRNAs with 80 pmol of RNP in HEK293T (**D**) cells or hESCs (**E**). Bars show averages (±SD) of at least three biological replicates.

It has previously been shown that crRNA and tracrRNA can be fused with a GAAA tetraloop or other linkers to yield a single guide RNA (sgRNA) with enhanced efficacy. Given the possibility that repeat:antirepeat interactions could affect efficacy, we explored the pairing between the repeat and anti-repeat of crRNA and tracrRNA. We designed and synthesized GC-rich crRNAs (**hiGC C1-C4**) and tracrRNAs (**hiGC T1-T4**) to improve pairing between crRNA and tracrRNA (Table S1). All of the modified RNAs outperformed *in vitro*-transcribed sgRNA as well as synthetic, unmodified dual RNAs (Figure S6). Furthermore, at lower concentrations, **hiGC**-**C1** exhibit increased potency relative to non-optimized versions of unmodified or modified RNAs (Figure S6). However, this trend does not hold true in *HTT*-**hiGC C1** (Figure S6). Therefore, these mutant sequences may be superior to wild-type sequences in a guide-sequence-specific manner.

In summary, we used a structure-guided approach combined with systematic addition of modifications to identify heavily- or fully-modified crRNAs and tracrRNAs that direct SpyCas9 genome editing in human cells. Two pairs of crRNA:tracrRNA stand out as particularly promising. First, **C20**:**T2** is our most potent combination, and both RNAs contain ribose substitutions at >80% of their nts. Furthermore, **C20** contains at least one chemical modification (2’-OMe, 2’-F or PS) at every single position. The **C20**:**T2** combination is more potent than its unmodified crRNA:tracrRNA counterpart when tested in human cells. Second, although the **C21**:**T8** combination exhibits reduced potency in human cells, its significant functionality is still noteworthy because it is completely devoid of ribose sugars. This will greatly ease chemical synthesis, enhance *in vivo* stability, and provide a springboard toward additional improvements (such as terminally appended chemical functionalities) that facilitate delivery and efficacy during clinical applications of genome editing.

## ASSOCIATED CONTENT

### Supporting Information

Supporting information (Materials and Methods, supplementary results, supplementary figures and tables) is available with the online version of this manuscript.

## AUTHOR INFORMATION

### Author Contributions

All authors participated in crRNA and tracrRNA design; A.M., M.R.H., A.J.D., and D.E. synthesized and purified crRNAs and tracrRNAs; A.M. expressed and purified recombinant SpyCas9; A.M. and E.H. conducted cellular genome editing experiments; A.M., J.F.A., J.K.W. and E.J.S wrote the manuscript; and all authors edited the manuscript.

### Notes

The authors declare the following competing financial interest(s): a patent application has been filed by the University of Massachusetts Medical School describing the inventions reported herein, with the authors as inventors. E. J. S. is a co-founder and Scientific Advisory Board member of Intellia Therapeutics.

## ACKNOWLEDGMENTS

The authors acknowledge partial support from the CHDI Foundation Research Contract A-10199 to M.H.B. We would also like to thank Nadia Amrani (Sontheimer lab) for her assistance with culturing hESCs. We are grateful to Scot Wolfe, Wen Xue, and members of their labs for materials and advice, and to all members of the Sontheimer, Watts, Khvorova, and Brodsky labs for helpful discussions.

