## Supplementary Materials for "Heavily and Fully Modified RNAs Guide Efficient SpyCas9-Mediated Genome Editing"

### METHODS

**Synthesis of Oligonucleotides.** CRISPR guides were synthesized at 1 µmole scale on an Applied Biosystems 394 DNA synthesizer. BTT (0.25 M in acetonitrile, ChemGenes) was used as activator. 0.05 M iodine in pyridine:water (9:1) (TEDIA) was used as oxidizer. DDTT (0.1 M, ChemGenes) was used as sulfurizing agent. 3% TCA in DCM (TEDIA) was used as deblock solution. Oligonucleotides were grown on 1000 Å CPG functionalized with Unylinker (~42 µmol/g). RNA and 2'-OMe phosphoramidites (ChemGenes) were dissolved in acetonitrile to 0.15 M; the coupling time was 10 min for each base. The nucleobases were deprotected with a 3:1 $NH_4OH$:EtOH solution for 48 hours at room temperature. Deprotection of the TBDMS group was achieved with DMSO:$NEt_3$•3HF (4:1) solution (500 µL) at 65 °C for 3 hours. RNA oligonucleotides were then recovered by precipitation in 3M NaOAc (25 µL) and n-BuOH (1 mL), and the pellet was washed with cold 70% EtOH and resuspended in 1 mL RNase-free water.

Purification of oligonucleotides were carried out by high performance liquid chromatography using a 1260 infinity system with an Agilent PL-SAX 1000 Å column (150 x 7.5 mm, 8 µm). Buffer A: 30% acetonitrile in water; Buffer B: 30% acetonitrile in 1M $NaClO_{4\,(aq)}$. Excess salt was removed with a Sephadex Nap-10 column.

Oligonucleotides were analyzed on an Agilent 6530 Q-TOF LC/MS system with electrospray ionization and time of flight ion separation in negative ionization mode. The data were analyzed using Agilent Mass Hunter software. Buffer A: 100mM hexafluoroisopropanol with 9mM triethylamine in water; Buffer B: 100mM hexafluoroisopropanol with 9 mM trimethylamine in methanol.

**Cell Culture.** A human HEK293T stable cell line expressing the traffic light reporter system was kindly provided by Wen Xue's lab in the RNA Therapeutics Institute at UMass Medical School. These cells were cultured in Dulbecco-modified Eagle's Minimum Essential Medium (DMEM; Life Technologies). DMEM was also supplemented with 10 % Fetal Bovine Serum (FBS; Sigma). HEK293T cells were obtained from ATCC and cultured in the same conditions. H1 human embryonic stem cells were cultured using feeder-free mTeSR medium (STEMCELL). Cells were grown in a humidified 37°C, 5% $CO_2$ incubator.

**Expression and Purification of 3xNLS-SpyCas9.** The pMCSG7 vector expressing the Cas9 from *Streptococcus pyogenes* was kindly provided by Dr. Scot Wolfe's lab. In this construct, the Cas9 also contains three nuclear localization signals (NLSs). Rosetta DE3 strain of *Escherichia coli* was transformed with the 3xNLS-SpyCas9 construct. For expression and purification of 3xNLS-SpyCas9, a previously described protocol was used.[1] The bacterial culture was grown at 37°C until an OD600 of 0.6 was reached.

Then, the bacterial culture was cooled to 18 °C, and 1 mM Isopropyl β-D-1-thiogalactopyranoside (IPTG; Sigma) was added to induce protein expression. Cells were grown overnight for 16-20 hrs.

The bacterial cells were harvested and resuspended in Lysis Buffer [50 mM Tris-HCl (pH 8.0), 5 mM imidazole]. 10 μg/mL of Lysozyme (Sigma) was then added to the mixture and incubated for 30 minutes at 4°C. This was followed by the addition of 1x HALT Protease Inhibitor Cocktail (ThermoFisher). The bacterial cells were then sonicated and centrifuged for 30 minutes at 18,000 rpm. The supernatant was then subjected to Nickel affinity chromatography as described by ref. [1]. The elution fractions containing the SpyCas9 were then further purified using cation exchange chromatography using a 5 mL HiTrap S HP column (GE). This was followed by a final round of purification by size-exclusion chromatography using a Superdex-200 column (GE). The purified protein was concentrated and flash frozen for subsequent use.

**Nucleofections of HEK293T cells.** The HEK293T cells were nucleofected using the Neon transfection system (ThermoFisher) according to the manufacturer's protocol. Briefly, 20-100 picomoles of 3xNLS-SpyCas9 was mixed with 25-125 picomoles of crRNA:tracrRNA in buffer R (ThermoFisher) and incubated at room temperature for 20-30 minutes. This Cas9 RNP complex was then mixed with approximately 100,000 cells which were already resuspended in buffer R. This mixture was nucleofected with a 10 μL Neon tip and then plated in 24-well plates containing 500 μL of DMEM and 10% FBS. The cells were stored in a humidified 37°C and 5% $CO_2$ incubator for 2-3 days.

**Flow Cytometry.** The nucleofected HEK293T cells were analyzed on MACSQuant® VYB from Miltenyi Biotec. For mCherry detection, the yellow laser (561 nm) was used for excitation and 615/20 nm filter used to detect emission. At least 20,000 events were recorded and the subsequent analysis was performed using FlowJo® v10.4.1. Cells were first sorted based on forward and side scattering (FSC-A vs SSC-A) to eliminate debris (**Figure S7**). Then cells were gated using FSC-A and FSC-H to select single cells. Finally, mCherry signal was used to select for mCherry-expressing cells. The percent of cells expressing mCherry was calculated and reported in this study as a measure of Cas9-based genome editing.

**Indel analysis by TIDE.** The genomic DNA from HEK293T cells was harvested using DNeasy Blood and Tissue kit (Qiagen) as recommended by the manufacturer. Approximately 50 ng of genomic DNA was used to PCR-amplify a ~700 bp fragment that was subsequently purified using a QIAquick PCR Purification kit (Qiagen). The PCR fragment was then sequenced by Sanger sequencing and the trace files were subjected to indel analysis using the TIDE web tool.[2]

***In vitro* DNA Cleavage Assays.** The traffic light reporter plasmid (Addgene: 31482) was linearized with restriction enzyme EcoRI (N.E.B.) for 1 hour, 37° C in NEB buffer 3, followed by heat inactivation for 20 minutes at 65°. For the Cas9 digest, 200 ng of linearized plasmid DNA was added to pre-formed RNP complexes (8 pmol or 0.8 pmol) and incubated for 1 hour at 37° in 25 μl NEB buffer 3. Cut DNA was purified using Zymo DNA purification columns and separated on a 1 % agarose gel run at 100 V. Relative intensities of full length and Cas9-cut DNA fragments were determined using Image J software.

**Serum Stability Assays.** A 10 μM Cas9 RNP complex was first assembled in cleavage buffer [20 mM HEPES (pH 7.5), 250 mM KCl and 10 mM $MgCl_2$]. Then 2 μM Cas9 RNP was incubated with 8% FBS in a 50 μL reaction at 37°C. Then at time points of 0 hr, 1 hr and 20 hrs, 10 μLs of the reaction mixture was treated with Proteinase K and then 10 uL of quench buffer (90% formamide and 25 mM EDTA) was added to the solution. The reaction mixture was resolved on a 10% denaturing polyacrylamide gel containing 6 M Urea. The gel was stained with SYBR Safe and visualized on Typhoon FLA imager.

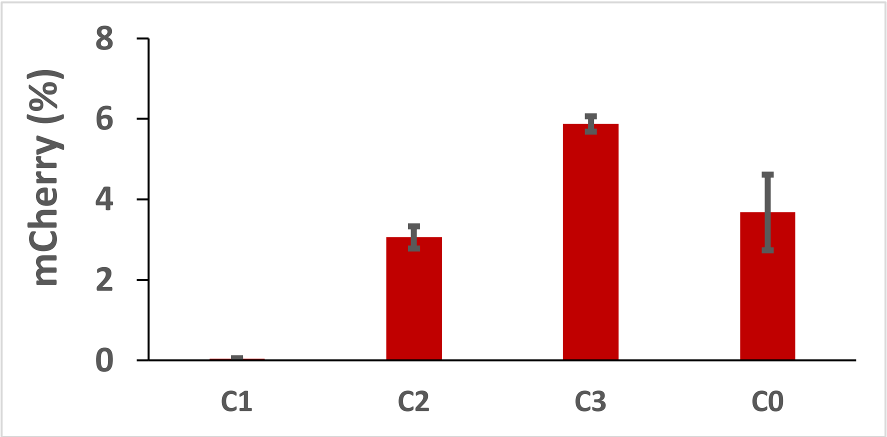

**Figure S1:** HEK293T-TLR cells were transfected with Lipofectamine CRISPRMAX Cas9 Transfection Reagent, and analyzed by flow cytometry for mCherry-positive cells.

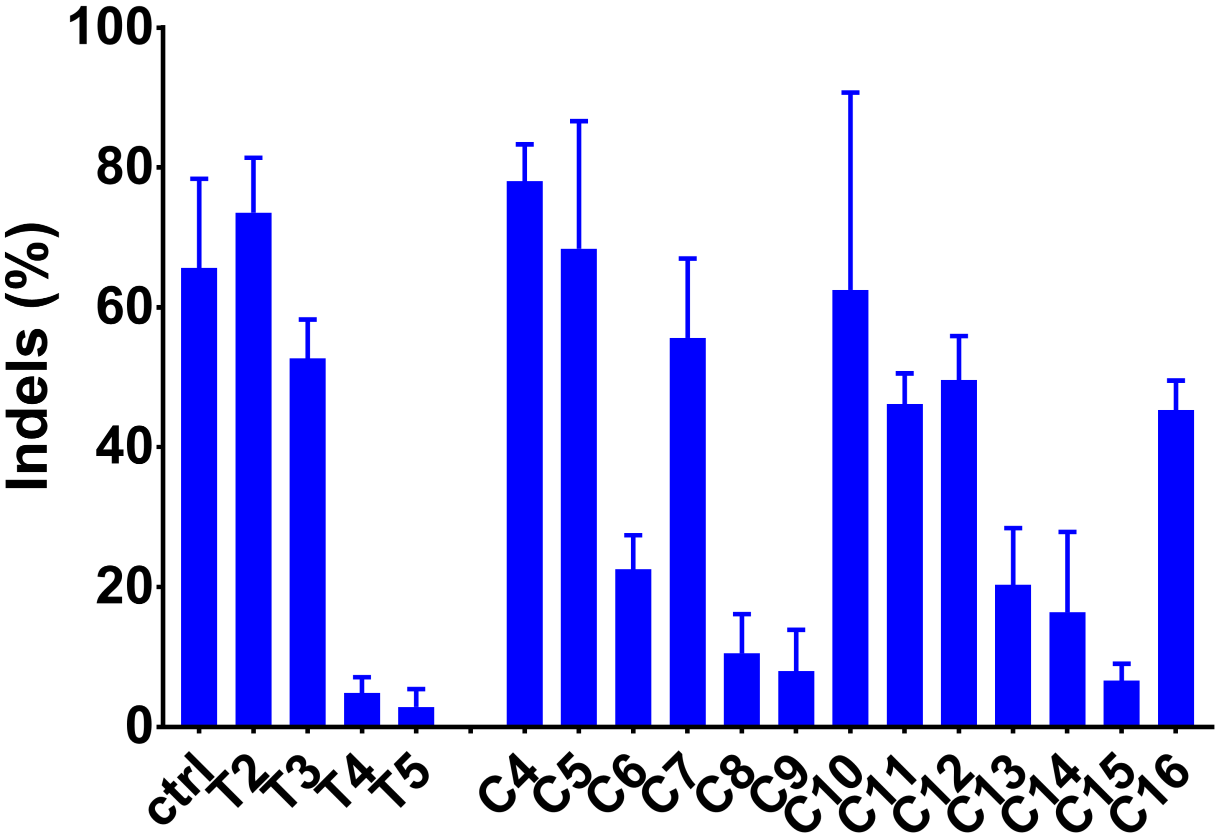

**Figure S2:** Overall editing efficiencies of Cas9 loaded with different modified RNAs. The chemically modified crRNAs **C4-C16** and tracrRNAs **T2-T5** were nucleofected into HEK293T-TLR cells and the resulting indel rate was determined using TIDE analysis. The modified crRNAs and tracrRNAs were tested against IDT purchased corresponding RNAs. The ctrl refers to IDT purchased crRNA:tracrRNA pair. Error bars represent standard deviations resulting from 3-5 biological replicates.

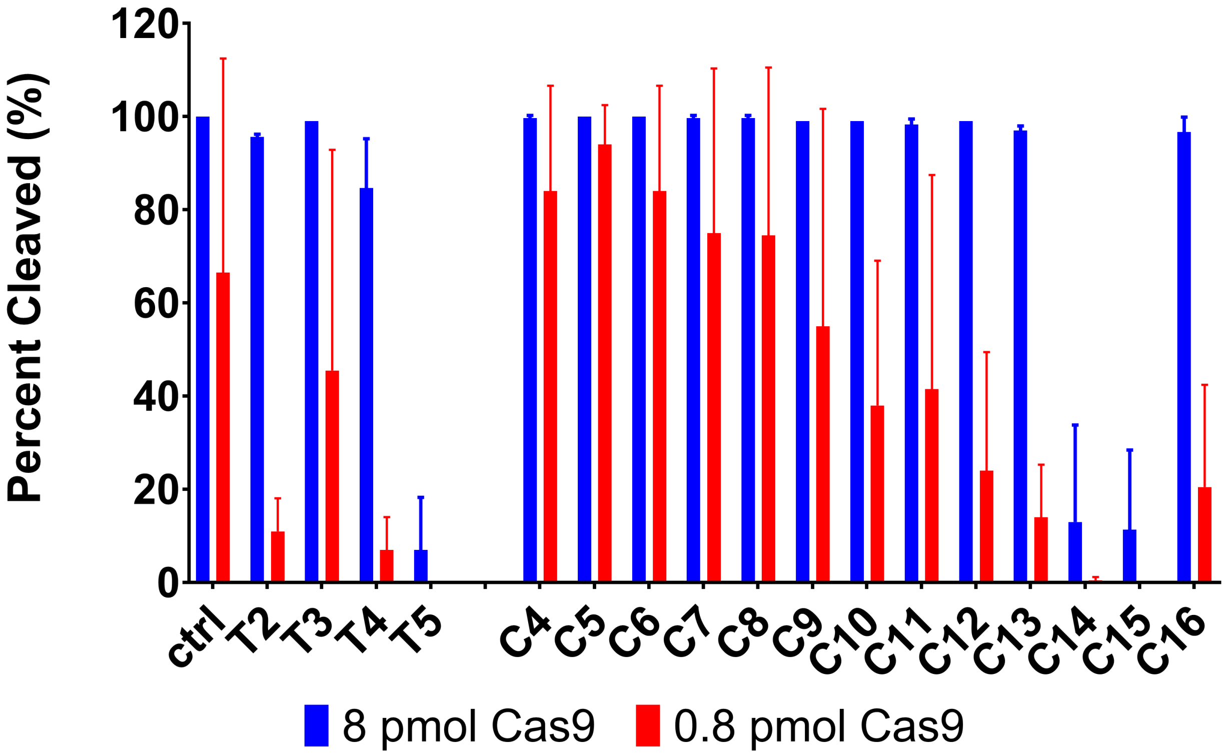

**Figure S3:** *In vitro* DNA cleavage assays to determine the functionality of modified RNAs. DNA cleavage assays were performed using saturating (8 pmols) and sub-saturating (0.8 pmols) amounts of Cas9 RNP complex. The modified crRNAs and tracrRNAs were tested against IDT purchased corresponding RNAs. The ctrl refers to IDT purchased crRNA:tracrRNA pair. Error bars represent standard deviations resulting from at least two replicates.

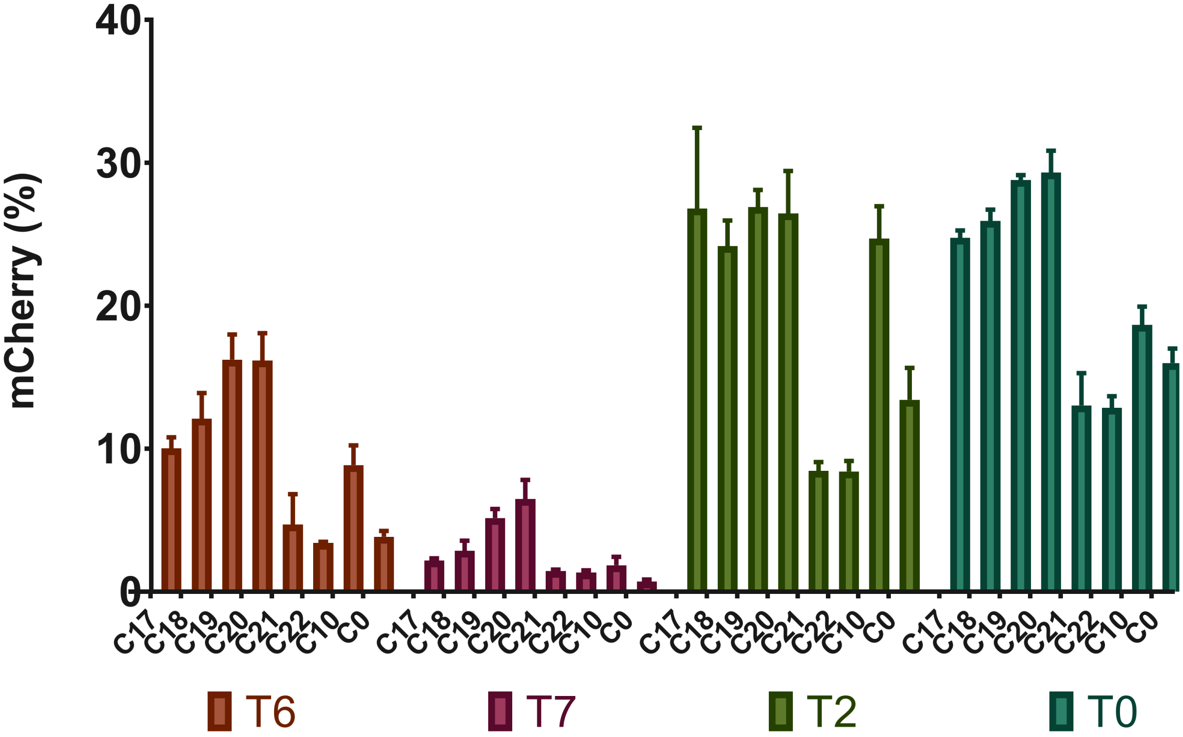

**Figure S4:** Comparison of synthetic crRNAs and tracrRNAs using 3 picomoles of Cas9 RNP (compare to **Figure 3** in the main text).

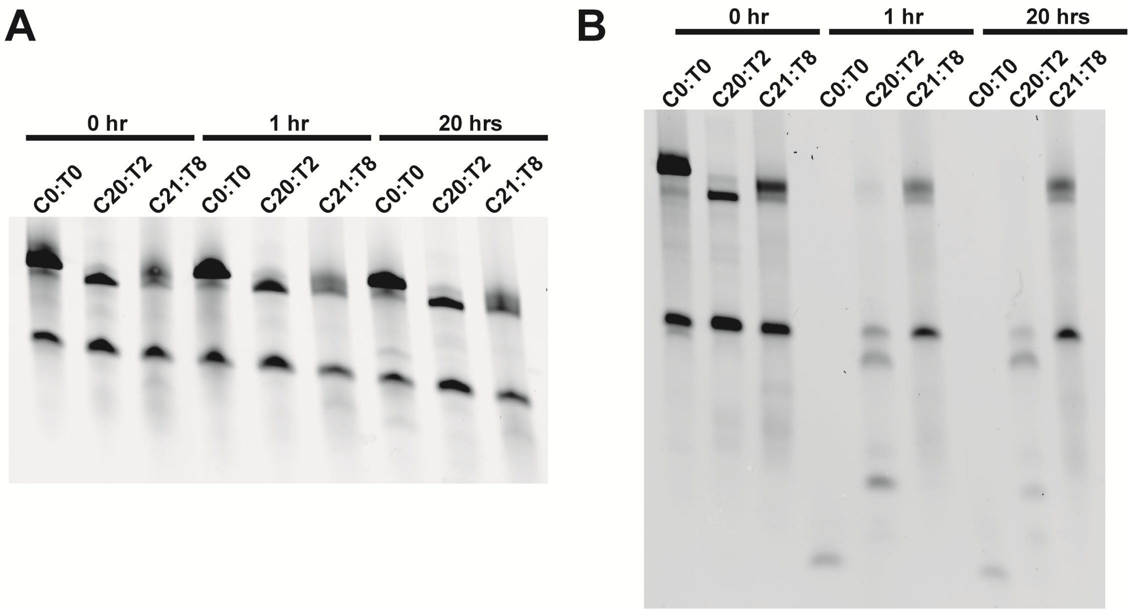

**Figure S5:** Serum stability of crRNAs C21, C0, C20 and tracrRNAs T0, T2, T8. The indicated crRNA:tracrRNA combinations were used to make Cas9 RNP complex that was then incubated with cleavage buffer (**A**) or 8% FBS (**B**) for 0, 1 and 20 hrs. The reactions were then treated with Proteinase K and then resolved on a 10% denaturing polyacrylamide gel. The gels were stained with SYBR Safe dye and then imaged on Typhoon FLA imager. The upper band in all lanes corresponds to tracrRNA and lower band corresponds to crRNA.

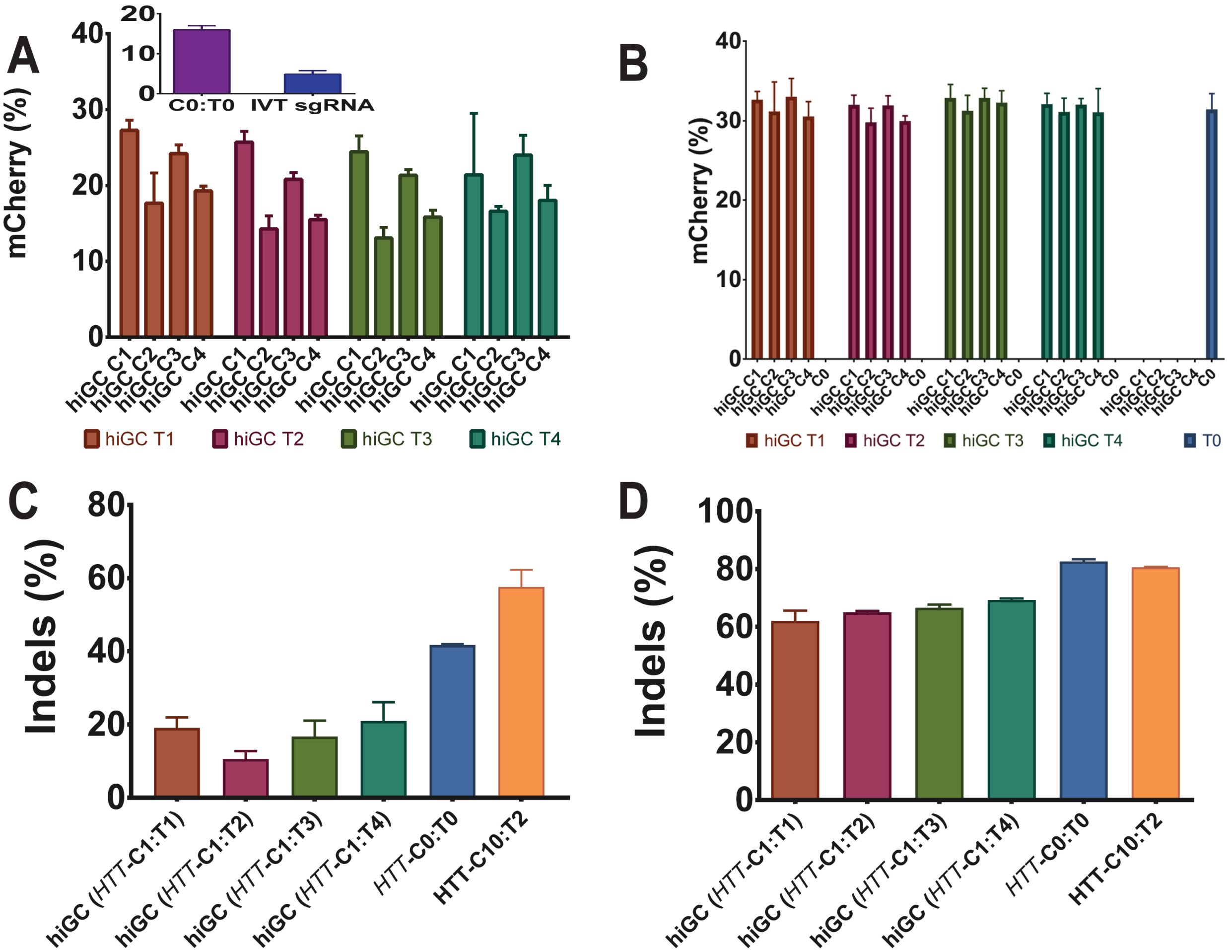

**Figure S6:** Comparison of sequence-optimized crRNA and tracrRNA series. HEK293T cells were nucleofected with 3 pmoles (**A & C**) or 20 pmoles (**B & D**) of Cas9 RNP. Two endogenous sites were targeted in HEK293T cells: the TLR site (**A & B**) and the genomic *HTT* site (**C & D**). **The** indel efficiency was determined using TIDE, whereas the fractions of mCherry-positive cells were measured using flow cytometry. *In vitro* synthesized (IVT) sgRNA was also included as a control in **A**. Error bars represent ±SD.

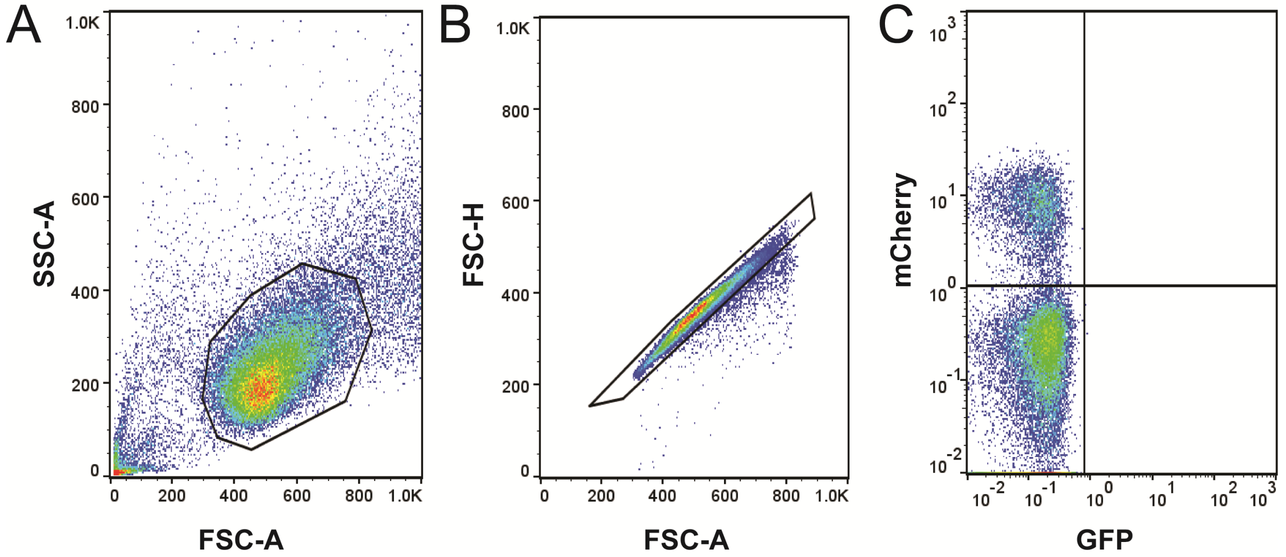

**Figure S7:** Flow cytometry analysis of HEK293T-TLR cells after nucleofection. The HEK293T cells were first gated based on forward and side scattering to remove cell debris (**A**), then gated to select single cells (**B**), and then finally gated to select mCherry-positive cells (**C**). The mCherry-positive cells are in the top left quadrant in **C**. The quadrant gate was based on the "no sgRNA" control in every experiment.

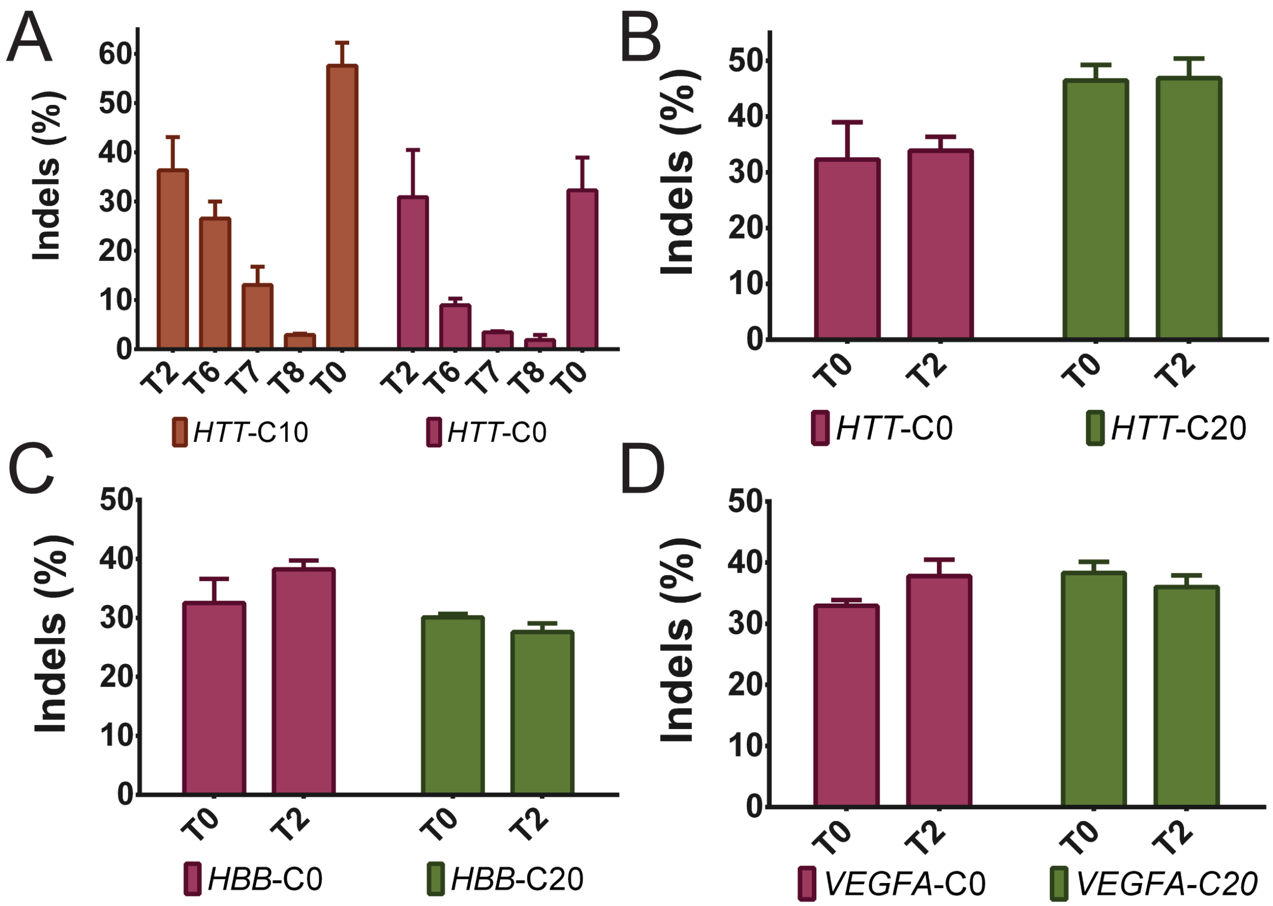

**Figure S8**. The crRNA designs **C10, C20** and **C21** tested with the indicated tracrRNAs using 3 pmoles of Cas9-RNP in HEK293T cells (compare to Figure 4 in the main text). Mean values from triplicate experiments (± SD) are shown. Three endogenous target sites were tested, namely *HTT, HBB* and *VEGFA*.

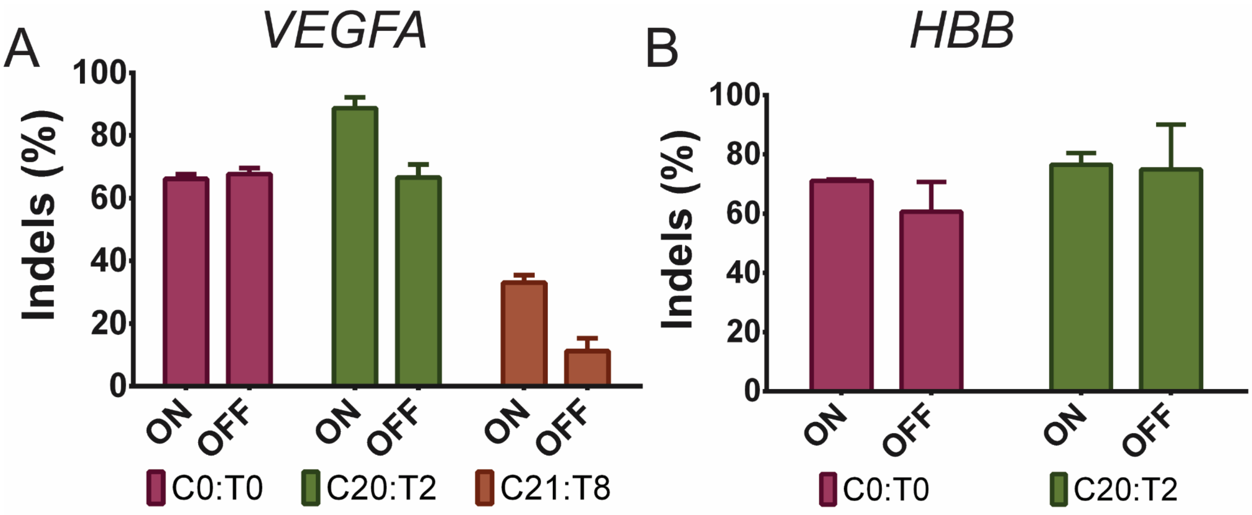

**Figure S9.** Effects of modified RNAs on on-target vs. off-target editing for target sites *VEGFA* and *HBB.*

**Table S1:** All the crRNAs and tracrRNAs synthesized for this study.

KEY: N = RNA, N = 2'-O-methyl RNA, N = 2'-fluoro RNA
N = 3' phosphorothioate

| Name | Sequence | Extinction Coefficient | MW (Observed/ Calculated) |
|---|---|---|---|
| **C1** | GGUGAGCUCUUAUUUGCGUAGUUUUAGAGCUAUGCU | 352710 | 11942.8/ 11942.8 |
| **C2** | GGUGAGCUCUUAUUUGCGUAGUUUUAGAGCUAUGCU | 352710 | 11739.6/ 11740.3 |
| **C3** | GGUGAGCUCUUAUUUGCGUAGUUUUAGAGCUAUGCU | 352710 | 11788.5/ 11788.5 |
| **C4** | GGUGAGCUCUUAUUUGCGUAGUUUUAGAGCUAUGCU | 352710 | 11888.6/ 11888.7 |
| **C5** | GGUGAGCUCUUAUUUGCGUAGUUUUAGAGCUAUGCU | 352710 | 11832.6/ 11832.6 |
| **C6** | GGUGAGCUCUUAUUUGCGUAGUUUUAGAGCUAUGCU | 352710 | 11832.6/ 11832.6 |
| **C7** | GGUGAGCUCUUAUUUGCGUAGUUUUAGAGCUAUGCU | 352710 | 11916.7/ 11916.8 |
| **C8** | GGUGAGCUCUUAUUUGCGUAGUUUUAGAGCUAUGCU | 352710 | 11916.6/ 11916.8 |
| **C9** | GGUGAGCUCUUAUUUGCGUAGUUUUAGAGCUAUGCU | 352710 | 12012.5/ 12013.2 |
| **C10** | GGUGAGCUCUUAUUUGCGUAGUUUUAGAGCUAUGCU | 352710 | 12024.5/ 12025.1 |
| **C11** | GGUGAGCUCUUAUUUGCGUAGUUUUAGAGCUAUGCU | 352710 | 12052.4/ 12053.3 |
| **C12** | Cy3-GGUGAGCUCUUAUUUGCGUAGUUUUAGAGCUAUGCU | 357610 | 12654.8/ 12655.5 |
| **C13** | Cy3-GGUGAGCUCUUAUUUGCGUAGUUUUAGAGCUAUGCU | 357610 | 12682.8/ 12683.6 |
| **C14** | Cy3-GGUGAGCUCUUAUUUGCGUAGUUUUAGAGCUAUGCUAAA-TegChol | 399190 | 14488.5/ 14489.3 |
| **C15** | Cy3-GGUGAGCUCUUAUUUGCGUAGUUUUAGAGCUAUGCUAAA-TegChol | 399190 | 14516.4/ 14517.5 |
| **C16** | Cy3-GGUGAGCUCUUAUUUGCGUAGUUUUAGAGCUAUGCUAAA-GalNAc | 399190 | 15522.2/ 15520.2 |
| **C17** | GGUGAGCUCUUAUUUGCGUAGUUUUAGAGCUAUGCU | 352710 | 12027.4/ 12027.2 |
| **C18** | GGUGAGCUCUUAUUUGCGUAGUUUUAGAGCUAUGCU | 352710 | 12026.4/ 12027.2 |
| **C19** | GGUGAGCUCUUAUUUGCGUAGUUUUAGAGCUAUGCU | 352710 | 12029.4/ 12029.2 |
| **C20** | GGUGAGCUCUUAUUUGCGUAGUUUUAGAGCUAUGCU | 352710 | 12000.4/ 12001.1 |
| **C21** | GGUGAGCUCUUAUUUGCGUAGUUUUAGAGCUAUGCU | 352710 | 11916.6/ 11916.6 |
| **C22** | GGUGAGCUCUUAUUUGCGUAGUUUUAGAGCUAUGCU | 352710 | 11972.5/ 11972.9 |
| **hiGC C1** | GGUGAGCUCUUAUUUGCGUAGUUUUAGAGCGAGCGC | 353160 | 12077.5/ 12078.1 |
| **hiGC C2** | GGUGAGCUCUUAUUUGCGUAGUUUUAGAGCGAGCGC | 353160 | 12005.4/ 12005.9 |
| **hiGC C3** | GGUGAGCUCUUAUUUGCGUAGUUUUAGAGCGAGCGC | 353160 | 12041.4/ 12042 |
| **hiGC C4** | GGUGAGCUCUUAUUUGCGUAGUUUUAGAGCGAGCGC | 353160 | 12029.4/ 12030 |
| **T1** | AGCAUAGCAAGUUAAAAUAAGGCUAGUCCGUUAUCAACUUGAAAA AGUGGCACCGAGUCGGUGCUUU | 699750 | 22130.8/ 22130.2 |
| **T2** | AGCAUAGCAAGUUAAAAUAAGGCUAGUCCGUUAUCAACUUGAAAA AGUGGCACCGAGUCGGUGCUUU | 699750 | 22439.6/ 22438.8 |
| **T3** | AGCAUAGCAAGUUAAAAUAAGGCUAGUCC GUUAUCAACUUGAAAAAGUGGCACCGAGUCGGUGCUUU | 699750 | 22632.5/ 22631.6 |
| **T4** | AGCAUAGCAAGUUAAAAUAAGGCUAGUCCGUUAUCAACUUGAAAA AGUGGCACCGAGUCGGUGCUUU | 699750 | 22453.6/ 22452.8 |
| **T5** | AGCAUAGCAAGUUAAAAUAAGGCUAGUCCGUUAUCAACUUGAAAA AGUGGCACCGAGUCGGUGCUUUAAA-TegChol | 741330 | 24287.4/ 24286.7 |
| **T6** | AGCAUAGCAAGUUAAAAUAAGGCUAGUCCGUUAUCAACUUGAAAA AGUGGCACCGAGUCGGUGCUUU | 699750 | 22449.2/ 22448.7 |
| **T7** | AGCAUAGCAAGUUAAAAUAAGGCUAGUCCGUUAUCAACUUGAAAA AGUGGCACCGAGUCGGUGCUUU | 699750 | 22453.1/ 22452.7 |
| **T8** | AGCAUAGCAAGUUAAAAUAAGGCUAGUCCGUUAUCAACUUGAAAA AGUGGCACCGAGUCGGUGCUUU | 699750 | 22463.3/ 22462.7 |
| **hiGC T1** | GCGCUCGCAAGUUAAAAUAAGGCUAGUCCGUUAUCAACUUGAAAA AGUGGCACCGAGUCGGUGCUUU | 681480 | 22407.4/ 22406.7 |
| **hiGC T2** | GCGCUCGCAAGUUAAAAUAAGGCUAGUCCGUUAUCAACUUGAAAA AGUGGCACCGAGUCGGUGCUUU | 681480 | 22335/ 22334.5 |
| **hiGC T3** | GCGCUCGCAAGUUAAAAUAAGGCUAGUCCGUUAUCAACUUGAAAA AGUGGCACCGAGUCGGUGCUUU | 681480 | 22371.3/ 22370.6 |
| **hiGC T4** | GCGCUCGCAAGUUAAAAUAAGGCUAGUCCGUUAUCAACUUGAAAA AGUGGCACCGAGUCGGUGCUUU | 681480 | 22335/ 22334.5 |
| ***HTT*-C10** | UGAAGUGCACACAGUAGAUGGUUUUAGAGCUAUGCU | 372510 | 12092.5/12093.2 |
| ***HTT*-C20** | UGAAGUGCACACAGUAGAUGGUUUUAGAGCUAUGCU | 363200 | 12092.5/ 12093.2 |
| ***HTT*-C21** | UGAAGUGCACACAGUAGAUGGUUUUAGAGCUAUGCU | 359820 | 12090.6/12090.7 |
| ***HTT*-hiGC 1** | UGAAGUGCACACAGUAGAUGGUUUUAGAGCGAGCGC | 372960 | 12121.5/12122.2 |
| ***HTT* C0** | UGAAGUGCACACAGUAGAUGGUUUUAGAGCUAUGCU | 372510 | 11679.3/11672.3 |

| | | | |
|---|---|---|---|
| **C0** | GGUGAGCUCUUAUUUGCGUAGUUUUAGAGCUAUGCU | 352710 | 11563.3/11564.1 |
| **T0** | Edit-R tracrRNA (Dharmacon) | -- | -- |
| ***HBB*-C0** | CUUGCCCCACAGGGCAGUAAGUUUUAGAGCUAUGCU | 351450 | 11583.2 |
| ***HBB*-C20** | CUUGCCCCACAGGGCAGUAAGUUUUAGAGCUAUGCU | 363200 | 12003.4/ 12004.1 |
| **VEGFA-C0** | GGUGAGUGAGUGUGUGCGUGGUUUUAGAGCUAUGCU | IDT | purchased |
| **VEGFA-C20** | GGUGAGUGAGUGUGUGCGUGGUUUUAGAGCUAUGCU | 363200 | 12175.4/ 12175.1 |
| **VEGFA-C21** | GGUGAGUGAGUGUGUGCGUGGUUUUAGAGCUAUGCU | 372510 | 12008.6/ 12008.7 |

(1) Jinek, M.; Chylinski, K.; Fonfara, I.; Hauer, M.; Doudna, J. A.; Charpentier, E. *Science* 2012, *337*, 816.

(2) Brinkman, E. K.; Chen, T.; Amendola, M.; van Steensel, B. *Nucleic Acids Research* 2014, *42*, e168.
